# Compensatory hepatic adaptation accompanies permanent absence of intrahepatic biliary network due to YAP1 loss in liver progenitors

**DOI:** 10.1101/2020.10.21.349159

**Authors:** Laura M. Molina, Junjie Zhu, Qin Li, Tirthadipa Pradhan-Sundd, Khaled Sayed, Nathaniel Jenkins, Ravi Vats, Sungjin Ko, Shikai Hu, Minakshi Poddar, Sucha Singh, Junyan Tao, Prithu Sundd, Aatur Singhi, Simon Watkins, Xiaochao Ma, Panayiotis V. Benos, Andrew Feranchak, Kari Nejak-Bowen, Alan Watson, Aaron Bell, Satdarshan P. Monga

**Author notes:** **Corresponding Author:** Satdarshan P. Monga, M.D., Professor of Pathology & Medicine (Gastroenterology, Hepatology & Nutrition), University of Pittsburgh, School of Medicine and UPMC, 200 Lothrop Street S-422 BST, Pittsburgh, PA 15261.

## Abstract

YAP1 regulates cell plasticity during liver injury, regeneration and cancer, but its role in liver development is unknown. YAP1 activity was detected in biliary cells and in cells at the hepato-biliary bifurcation in single-cell RNA-sequencing analysis of developing livers. Hepatoblast deletion of *Yap1* led to no impairment in Notch-driven SOX9+ ductal plate formation, but prevented the formation of the abutting second layer of SOX9+ ductal cells, blocking the formation of a patent intrahepatic biliary tree. Intriguingly, the mice survived for 8 months with severe cholestatic injury and without any hepatocyte-to-biliary transdifferentiation. Ductular reaction in the perihilar region suggested extrahepatic biliary proliferation likely seeking the missing intrahepatic biliary network. Long-term survival of these mice occurred through hepatocyte adaptation via reduced metabolic and synthetic function including altered bile acid metabolism and transport. Overall, we show YAP1 as a key regulator of bile duct development while highlighting a profound adaptive capability of hepatocytes.

## Introduction

The biliary tree is a delicate branching network of ducts formed of cholangiocytes which transport bile from the liver to the intestines. Alagille syndrome is an autosomal dominant disorder arising from mutations in the *JAGGED1* or *NOTCH2* genes which causes multi-system malformations including impaired formation of bile ducts in embryonic development (Mitchell et al., 2018). According to a recent prospective study, only about 24% of children with bile duct paucity reach adulthood without a liver transplant, indicating the serious need for alternative therapies (Kamath et al., 2020). Interestingly, the penetrance of these mutations varies widely leading to variability in the extent of cholestasis and disease presentation, with some patients even showing spontaneous recovery. We lack an understanding of the disease modifiers and relevant biomarkers that can help stratify or distinguish these patients during a critical treatment window (Emerick et al., 1999; Lykavieris et al., 2001; Mitchell et al., 2018).

Yes-associated protein 1 (YAP1) is a transcriptional co-activator, and a mechanosensor that modulates cell differentiation, proliferation and survival among liver cells depending on the context (Patel et al., 2017). Studies have shown that YAP1 is important for bile duct development and homeostasis, although its exact role remains poorly understood (Lee et al., 2016; Pepe-Mooney et al., 2019; Yimlamai et al., 2014; Zhang et al., 2010). Albumin (Alb)-Cre mediated deletion of YAP1 in late stages of murine liver development led to bile duct paucity postnatally, causing unresolved cholestatic injury (Zhang et al., 2010). Similarly, activation of YAP1 through Alb-Cre mediated deletion of upstream negative regulators LATS1/2 resulted in abnormal overgrowth of ductular cells, and *in vitro*, facilitated hepatic progenitor differentiation into BECs (Lee et al., 2016). Other studies have shown YAP1 as a major driver of hepatoblastoma, a pediatric liver tumor, and can also dedifferentiate mature hepatocytes into hepatoblasts (HBs) in a murine model (Alder et al., 2014; Molina et al., 2019; Tao et al., 2014). Thus, the role of YAP1 in hepatobiliary differentiation remains ambiguous.

The hepatobiliary plasticity is being increasingly appreciated. Chronic injury to the bile ducts in rodents induced transdifferentiation of hepatocytes into cholangiocytes to promote repair (Michalopoulos et al., 2005; Okabe et al., 2016; Schaub et al., 2018; Sekiya and Suzuki, 2014; Yanger et al., 2013). Recently this was convincingly observed in an animal model of Alagille syndrome with liver-specific developmental ablation of Notch signaling and HNF6 (Schaub et al., 2018). Intriguingly, despite the total failure of intrahepatic bile duct formation, many of these mice recovered and survived long-term due to hepatocyte-derived *de novo* generation of bile ducts (Schaub et al., 2018). Phenotypic recovery over time has also been observed in some (but not all) murine models of Alagille syndrome (Andersson et al., 2018; Thakurdas et al., 2016). While YAP1 can promote the expression of biliary markers in mature hepatocytes (Yimlamai et al., 2014) and is a potential regulator for Notch signaling (Kim et al., 2017; Tschaharganeh et al., 2013), it is not known whether YAP1 activation is essential or dispensable for cells to adopt a biliary phenotype or to assemble into functional ductular structures.

In this study, we conclusively address the role of YAP1 during the earliest stages of embryonic liver development demonstrating its indispensable role in bile duct morphogenesis. The conditional loss of YAP1 in HBs led to a complete failure of intrahepatic biliary tree generation, reminiscent of Alagille syndrome, as demonstrated using a variety of functional studies and innovative three-dimensional (3D) imaging. Further, we characterize the compensatory metabolic and synthetic adaptations that allow mice with severe cholestatic injury to survive long-term.

## Results

### Loss of YAP1 in HBs during early embryonic development leads to absence of intrahepatic bile duct formation and severe chronic cholestatic injury

To assess the status of YAP1 activity in liver development, we analyzed single-cell sequencing data published by Yang L. *et al*, for the expression of YAP1 targets (Yang et al., 2017). This study had previously assessed the differentiation of HBs into hepatocytes and cholangiocytes from E10.5 to E17.5. We performed Louvain clustering and tSNE visualization and identified three clusters closely matching the cell identities assigned by Yang L. *et al*, clearly distinguishing the trajectory of differentiation over pseudotime of HBs (cluster a) into hepatocytes (cluster c) and cholangiocytes (cluster b) (Fig.S1A-B). When comparing gene expression in these clusters, we observed that the expression of canonical YAP1 targets *Ccn1* and *Ccn2* was notably increased in the cholangiocyte cluster over the pseudotime axis while remaining low in the HB and hepatocyte clusters (Fig.S1C-D). We also found that several cells from early time points E11.5 – E14.5 were classified differently by our methods in comparison to those previously published, and express both *Sox9* and *HNF4α*, suggesting they may be cells of intermediate differentiation (Fig.S1A-B). YAP1 target gene expression in these cells (cluster d) is somewhat higher than in most HBs but lower than most cholangiocytes (Fig.S1C), suggesting that YAP1 activity is increasing in these intermediate cells as they transition into cholangiocytes.

Previous studies have used *Alb-Cre* to delete YAP1 in the liver to elucidate its functions. However, in this model recombination occurs around E15-16 and is not completed until 4-6 weeks postnatally (Postic and Magnuson, 2000). Since the process of cholangiocyte differentiation from HBs begins earlier at around E11.5 (Su et al., 2017; Yang et al., 2017), at which time we also observe YAP1 activation, we targeted YAP1 in HBs at the earliest stages of development. Since *Foxa3* is activated in the foregut endoderm progenitors, including HBs, starting at E8.5 and achieving complete recombination around E12-13 (Lee et al., 2005; Tan et al., 2008), we bred *Foxa3-Cre* transgenic to YAP1-floxed mice and generated HB-YAP1 knockout (YAP1 KO) mice. IF imaging for YAP1 at E14.5 shows complete loss of YAP1 from HNF4α-labeled HBs (Fig.S1E).

Although we expected to observe a more widespread effect, the YAP1 KOs formed all foregut organs and were viable postnatally, surviving up to 8 months at which time they required euthanasia. Grossly, mice displayed stunted growth and were visibly jaundiced (Fig.1A), with significantly lower body weight than WT littermates, although their liver weights were comparable to WT mice, resulting in a dramatically increased liver weight to body weight ratio (Fig.1B-D). At postnatal day 21 (P21), YAP1 KO mice showed elevated alkaline phosphatase, total bilirubin and direct bilirubin levels, and significantly elevated aspartate and alanine aminotransferases (AST and ALT), indicating severe hepatocellular and cholestatic injury (Fig.1E-H). H&E staining displayed numerous biliary infarcts (Fig.S1F). IHC for YAP1 in adult KO mice showed persistent complete YAP1 loss in all liver epithelial cells, but retained YAP1 staining in endothelial cells and other non-parenchymal cells, as compared to WT which showed low levels of cytoplasmic YAP1 in most hepatocytes and strong nuclear YAP1 in mature bile ducts (Fig.1I, L). CK19 staining shows that while WT livers have well-formed bile ducts surrounding the portal veins throughout the liver, YAP1 KO mice demonstrated an absence of defined ductal structures except for a few ducts limited to the perihilar region and even fewer large ducts extending occasionally into the median hepatic lobe (Fig.1J, K, M). SOX9 staining in adult KO livers revealed unstructured clusters of SOX9+ cells around the portal vein, in contrast to well-formed SOX9+ patent bile ducts in WT (Fig.1G).

**Figure 1.**
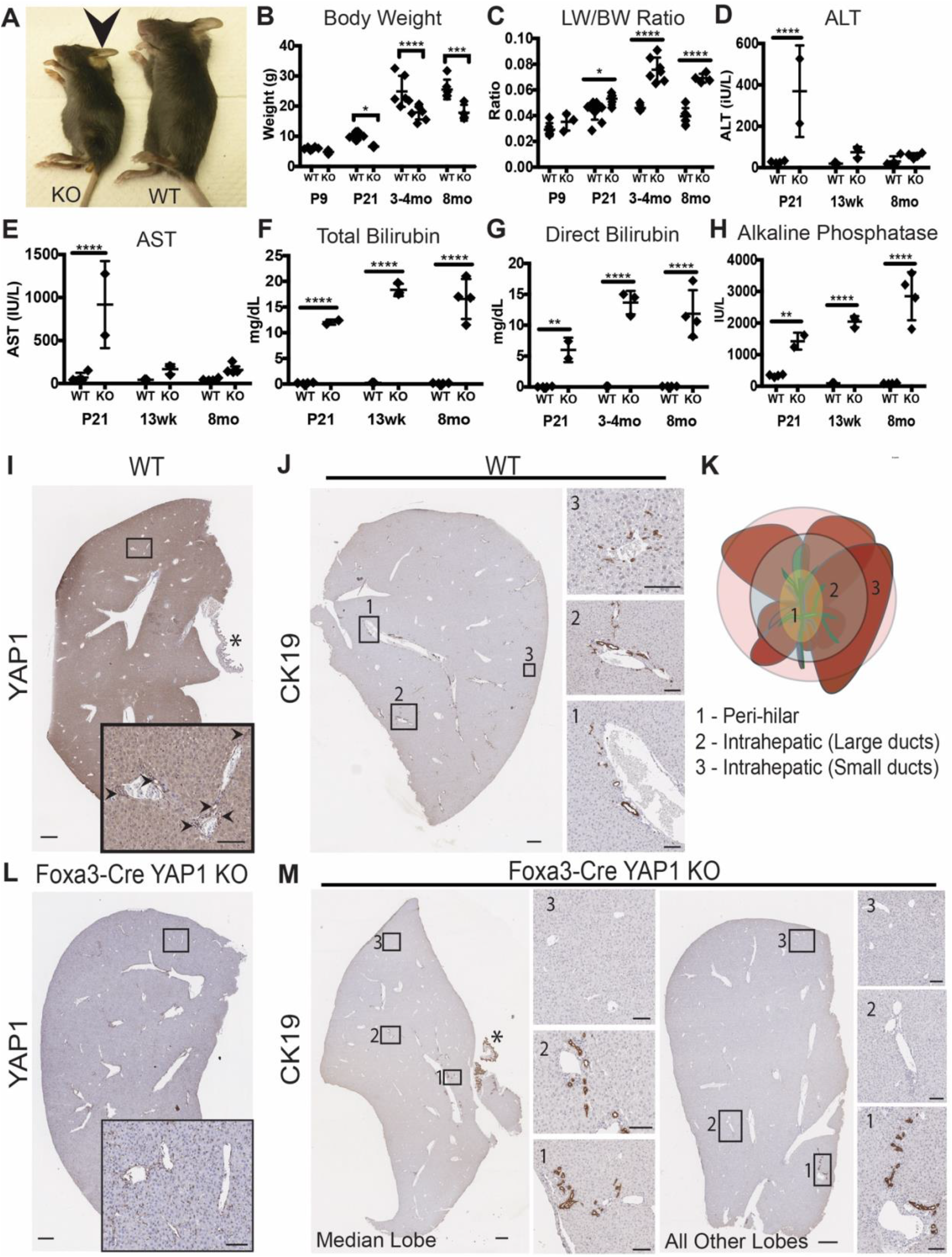
Loss of YAP1 in HBs leads to failure of intrahepatic bile duct formation. A. Gross image of WT and YAP1 KO mouse, arrow shows jaundiced ears. B. Body weight and C. Liver weight to body weight (LW/BW) ratio of WT and KO mice over time. Serum levels of D. alanine aminotransferase (ALT), E. aspartate aminotransferase (AST), F. total bilirubin, G. direct bilirubin, and H. alkaline phosphatase in WT and KO mice over time. Graphs show mean ± sd. Data were analyzed by 2-way ANOVA with Sidak multiple comparison test, n = 2-5 mice per group (* p<0.05, ** p < 0.01, *** p<0.001, **** p<0.0001). I. IHC for YAP1 in WT mice; arrows show nuclear YAP1 in bile ducts. J. CK19 marks bile ducts in the liver spanning three main regions described in panel K. L. IHC for YAP1 in KO mice. M. CK19 staining in various lobes of YAP1 KO mice. Asterisks mark the gallbladders. Scale bars are 500μm for whole lobes, 100μm for insets.

Surprisingly, YAP1 KO mice survived for over 8 months when they were euthanized due to progressive morbidity. Even though markers of cholestatic injury remained severely elevated throughout, AST and ALT returned to almost normal levels by 3 months of age, suggesting that these injured livers deployed some adaptive mechanisms to survive despite severe cholestasis (Fig.1E-H).

Thus overall, YAP1 KO mice exhibited significant failure to thrive and cholestatic injury, associated with persistent lack of intrahepatic bile ducts in the postnatal liver, resembling Alagille syndrome-like phenotype.

### YAP1 KO mice show no hepatocyte-driven biliary regeneration, but exhibit limited DR around hilum, arising from extrahepatic bile ducts

We next investigated repair mechanisms that may be allowing YAP1 KO to survive long-term. Previous studies have convincingly demonstrated the capacity of normal hepatocytes to transdifferentiate to regenerate *de novo* bile ducts in the setting of similar developmental biliary defects *in vivo* (Schaub et al., 2018). Hence, we first evaluated whether any similar evidence of biliary regeneration occurred over time in YAP1 KO. Intriguingly, we found no YAP1-negative bile ducts even up to 8 months. In fact, majority of the liver remained devoid of any intrahepatic ducts. To unequivocally demonstrate this, we visualized the 3D structure of the bile ducts in WT and YAP1 KO mice at P21 and 8 months using a novel tissue clearing protocol combined with ribbon scanning confocal microscopy, to achieve 3D IF staining of whole liver tissue (Fig.2A-D, Movie.S1). Staining with CK19 clearly delineated the hierarchical branching biliary network in P21 and 8-month-old WT mice, which was completely absent in YAP1 KO mice at both times except for a few ducts in the perihilar region connected to the extrahepatic biliary tree and gallbladder. Thus, we unambiguously demonstrate the gross absence of biliary regeneration in YAP1 KO mice.

**Figure 2.**
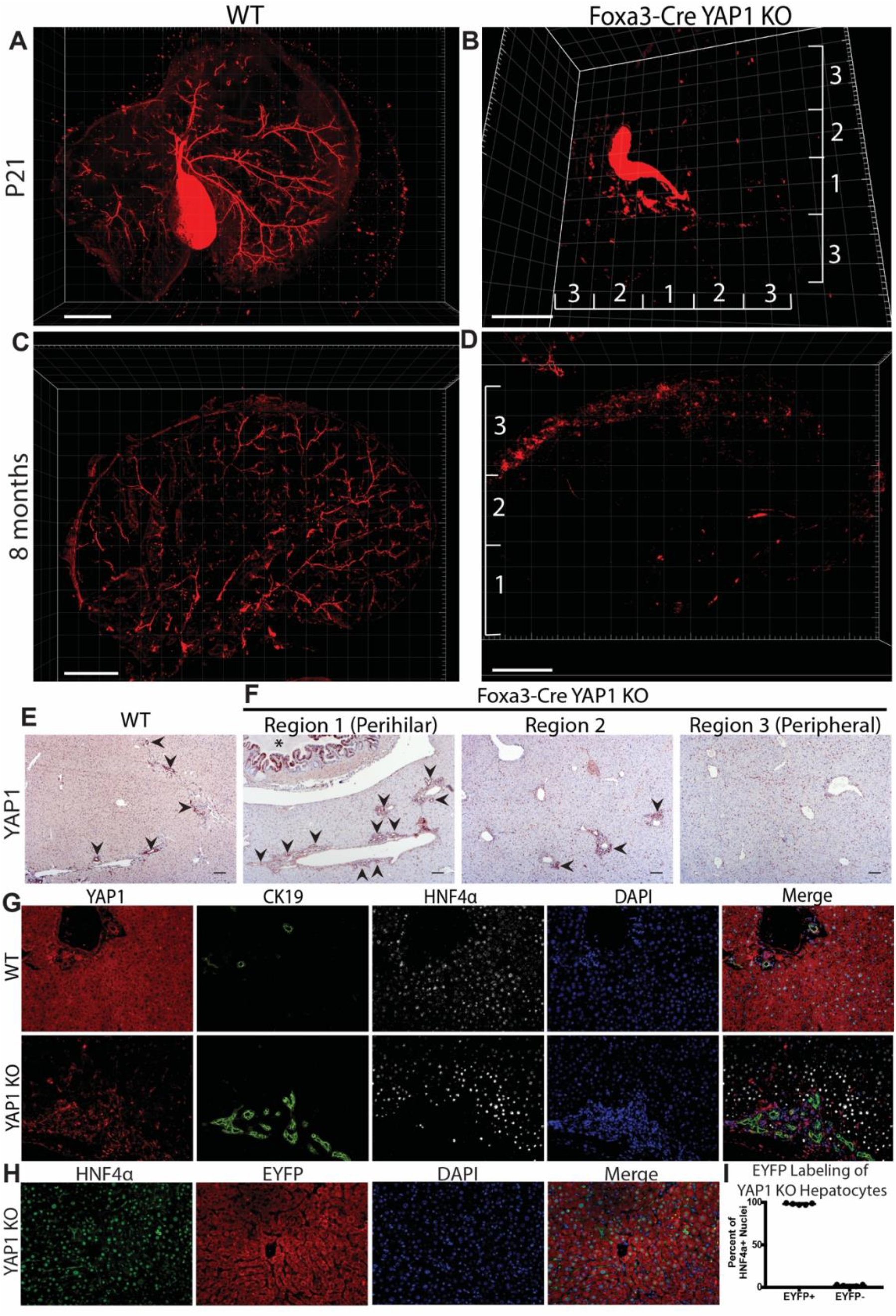
YAP1 KO mice show no long-term regeneration of bile ducts and no transdifferentiation of hepatocytes into cholangiocytes. IF for CK19 followed by ribbon-confocal scanning microscopy illustrate in 3D the mature biliary tree of WT mice at (A) P21 and (C) 8 months of age and show the absence of bile ducts in YAP1 KO mice at (B) P21 and (D) 8 months of age. Scale bars are 2mm. Regions 1, 2, and 3 refer approximately to the expected positions of perihilar, intrahepatic large ducts, and intrahepatic small ducts respectively. E. IHC for YAP1 in WT mice (arrows highlight bile ducts). F. IHC for YAP1 in KO mice (arrows highlight ductular reaction). Scale bars are 100μm. G. IF co-staining for YAP1, CK19, and HNF4α in WT and YAP1 KO mice (20x magnification). H. IF co-staining for HNF4α and EYFP in YAP1 KO mice (20x magnification). I. Quantification of EYFP-positive HNF4α-positive cells in adult YAP1 KO mice (mean ± sd, n=5 mice, representing the average of 3-5 20x fields per mouse).

Interestingly, we noticed ductular reaction (DR) *albeit* only in the perihilar region and particularly in the median lobe of the mouse liver. This DR consisted of YAP1positive ducts and associated with significant inflammation and fibrosis surrounding the largest portal veins (Fig.2E-F, Fig.S1H-I). To assess the source of DR, we wondered whether the YAP1-positive ducts could have arisen from YAP1-positive hepatocytes that may have escaped Cre-recombination and could have then transdifferentiated into BECs. However, no YAP1-positive hepatocytes were observed by IHC (Fig.1L). IF showed that while the DR in hilar region was YAP1-positive, the surrounding hepatocytes were YAP1-negative (Fig.2G). Using a ROSA-stop^fl/fl^-EYFP reporter, we verified that >99% of hepatocytes (HNF4α+) in YAP1 KO mice were EYFP-positive demonstrating successful Cre-recombination (Fig.2H-I). Thus, we rule out hepatocytes as a source of the DR, and also show that YAP1-negative hepatocytes are not capable of transdifferentiating into biliary cells validating the role of YAP1 in hepatocyte transdifferentiation (Fitamant et al., 2015; Yimlamai et al., 2014).

To further substantiate the functional absence of intrahepatic biliary tree and at the same time assess if the regional DR contributed functionally, we measured the bile flow rate by cannulating the common bile duct, which is present in both WTs and YAP1 KOs. While expected bile flow at the expected normal rate in an unstimulated setting was evident in WT mice, there was no detectable bile flow in YAP1 KO mice (Fig.3A). Altogether, we show lack of intrahepatic biliary tree and a DR from extrahepatic ducts restricted mostly to the perihilar region, which was insufficient to restore function through intrahepatic biliary repair in YAP1 KO mice.

**Figure 3.**
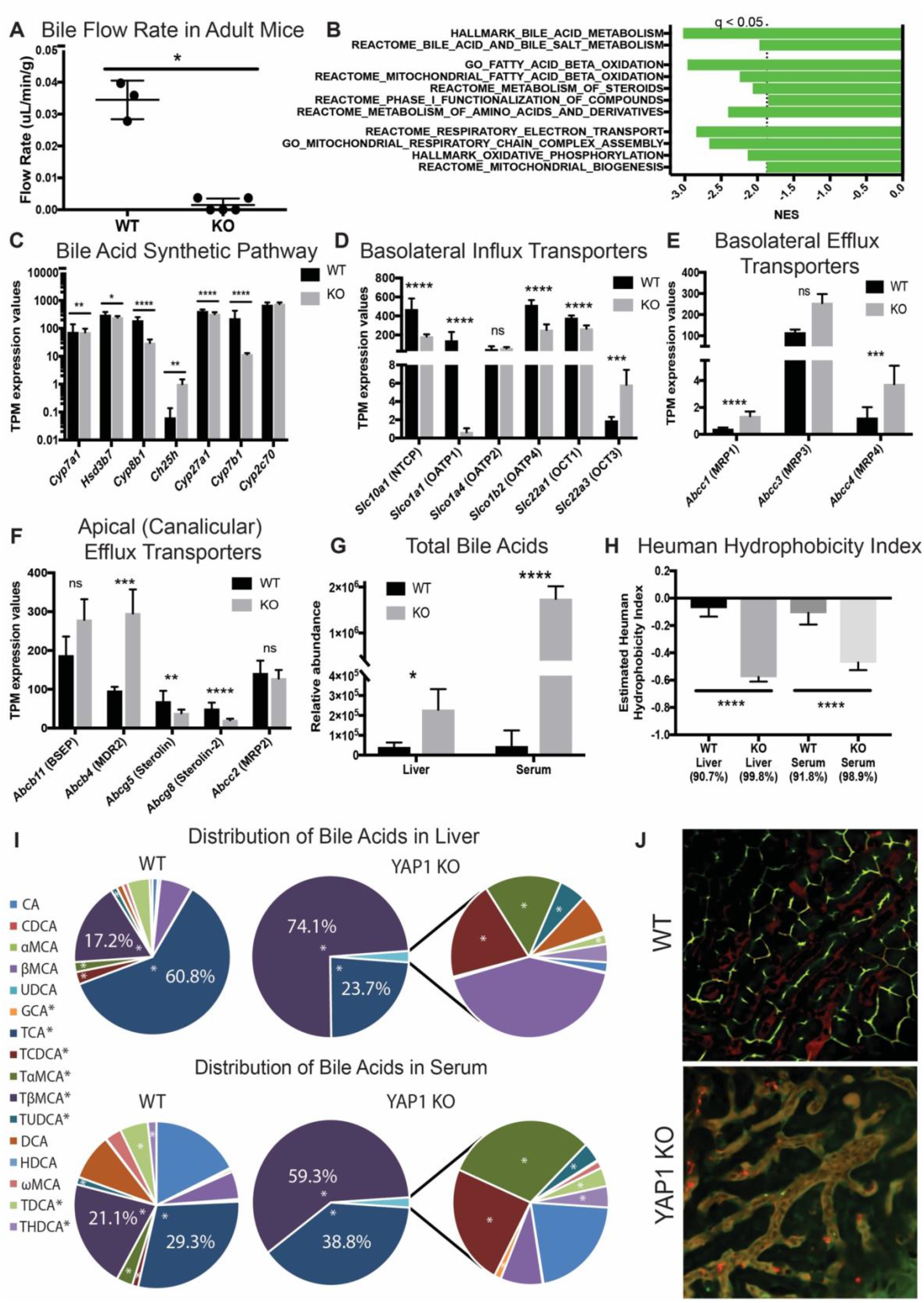
YAP1 KO mice adapt to chronic cholestasis by reducing bile acid toxicity and secreting them into the bloodstream. A. Cannulation of the common bile duct was used to measure baseline bile flow in adult WT and YAP1 KO mice (3-5 mice per group, two-tailed Mann-Whitney test, *p<0.05). B. GSEA revealed several metabolic pathways negatively enriched in YAP1 KO mice vs WT. C-F. RNA-seq analysis shows altered gene expression of genes related to bile acid synthesis and excretion (*q<0.05, **q<0.01, ***q<0.001, ****q<0.0001). G. Mass spectrometry was used to measure the abundance of bile acid species in liver tissue and serum from WT and YAP1 KO mice (n of 8 WT with 4 males and 4 females, and 7 KO with 4 males and 3 females; data show mean ± sd; 2-way ANOVA with Sidak multiple comparison test, * p<0.05, **** p<0.0001). H. The hydrophobicity index of the bile acid pool in liver and serum was calculated based on the Heuman index values for each bile acid species(Heuman, 1989) (mean ± sd, t-test, **** p<0.0001). Underneath we include the percentage of total bile acids used in each calculation as the index values for certain species are unavailable. I. The average distribution and abundance of murine bile acid species is shown for WT and YAP1 mice. Asterisks (*) refer to conjugated bile acids. J. Still shots taken from live movies (Supplemental Movie 2) from intravital microscopy showing the circulation of blood and bile in both WT and YAP1 KO mice.

### Gene expression changes in YAP1 KO livers indicate global metabolic and synthetic changes including in bile acid homeostasis as an adaptation to the absence of bile ducts

Since biliary reconstitution was not the basis of prolonged survival in the YAP1 KO, we posited that several adaptations to chronic cholestatic injury may be in play. We performed RNA-sequencing analysis on adult YAP1 KO mice and WT mice, using both males and females in each group for comparison. Principal component analysis showed WT and KO mice to be distinguished by the first principal component (Fig.S2A). Comparison of YAP1 KO and WT livers identified 2606 differentially expressed genes (FDR<0.05, abs(FC)>2). Pathway analysis using various common software algorithms – Ingenuity Pathway Analysis (Qiagen), Gene Set Enrichment Analysis(Subramanian et al., 2005), and Enrichr (Kuleshov et al., 2016), were performed and similar observations were evident. Both pathway analysis and transcription factor enrichment analysis highlighted robust activation of innate and adaptive immune responses in YAP1 KO mice, along with stellate cell activation and fibrosis (Fig.S2B-C, E-F). This is consistent with histological evidence of increased pericellular fibrosis and presence of inflammatory cell clusters in adult KO (Fig.S1H-I).

In addition, we observed an increase in pathways related to proliferation, cell cycling, and cancer alongside downregulation of mature metabolic and synthetic genes normally expressed in hepatocytes (Fig.S2D-F). We also observed a significant downregulation of broad metabolic and synthetic pathways related to fatty acid oxidation, oxidative phosphorylation, xenobiotic metabolism, and bile acid and sterol metabolism, all of which are hallmarks of mature liver function (Fig.3B). Decreases in fatty acid oxidation have been previously described in multiple cholestatic diseases in mice and patients and shown to be related to decreased PPARα activity, which was also supported in YAP1 KO by IPA and Enrichr analysis (Fig.S2E-F) (Katzenellenbogen et al., 2007; Pablo Arab et al., 2017; Zhao et al., 2017). Decreased gene expression related to bile acid and sterol metabolism suggested inactivation of FXR, RXR, and LXR transcription factors (Fig.S2F). Likewise, expression of most enzymes involved in the classic and alternative bile acid synthesis pathways were downregulated in YAP1 KO livers (Fig.3C). Interestingly, expression of most apical and basolateral transporters regulating bile acid transport in hepatocytes was altered to favor increased secretion back into the sinusoids rather than the hepatocyte apical canaliculi (Fig.3D-F).

To assess the functional consequences of these gene expression changes, we performed quantitative and qualitative analysis of bile acids in YAP1 KO mice compared to WT in both liver tissue and serum. The total quantity of bile acids was dramatically increased in the liver (~6x) and in the serum (~40x) in YAP1 KO mice (Fig.3G) suggesting the general decrease in bile acid synthetic enzyme expression to be compensatory. Next, in the KOs, the diversity of species normally found in WT mice, was overwhelmingly shifted in favor of primary conjugated bile, particularly taurocholic acid (TCA) and taurobetamurocholic acid (TβMCA), which individually were increased by almost 1000-fold in the serum, while the less soluble primary unconjugated bile acids were significantly decreased in the liver tissue (Fig.3I, Fig.S3A-L). Secondary bile acids derived from bacterial digestion of primary bile acids in the gut were also significantly decreased in the liver and serum of KO mice (Fig.S3A-L). Using previously published data on bile acid hydrophobicity (Heuman, 1989), we show that the bile acid pool in both liver and serum of YAP1 KO mice exhibited significantly lower hydrophobicity indices compared to WT, indicating a shift toward more soluble bile acid species to reduce their toxicity and facilitate their secretion into the blood (Fig.3H).

We next used intravital microscopy to visualize the flow of bile from hepatocytes into the canaliculi in WT and KO (Fig.3J, Movie.S2). In WT mice, carboxy-fluorescein-di-acetate (CFDA) injected into the bloodstream was taken up by hepatocytes, metabolized into carboxyfluorescein (CF) to fluoresce green, and exported into the hepatocyte canaliculi, thus providing a clear view of the tightly sealed canalicular network completely segregated from the blood (dyed red using TXR Dextran) flow in sinusoids. In contrast, in YAP1 KO livers, none of the CF entered the canaliculi which could not be visualized. There was notable mixing of blood and bile as shown by the yellow color in sinusoids suggesting hepatocyte-metabolized CF transporting back into the blood (Fig.3J, Movie.S2). Thus overall, these transcriptional and functional adaptations reflect a concerted effort to remove bile acids from liver by exporting them into the serum while also reducing their toxicity through conjugation, in an attempt to limit hepatocellular injury.

### YAP1 loss does not impair ductal plate formation but prevents formation of second layer of ductal cells thus impairing bile duct morphogenesis

Although YAP1 has previously been suggested to play a role in the formation of bile ducts in embryonic development, a conclusive evidence and underlying mechanism has remained elusive. To investigate, we closely examined previously described stages of intrahepatic bile duct formation in YAP1 KO livers. SOX9-positive cells representing the initial formation of the ductal plate, were evident around the portal veins at E14.5 and E16.5, similarly in the WT and YAP1 KO (Fig.4A-B). HES1, a downstream target of Notch signaling, was also similarly expressed in the ductal plate cells of the WT and KO embryonic livers (Fig.S4C), suggesting initial activation of Notch signaling in putative BECs is not affected by YAP1 loss.

**Figure 4.**
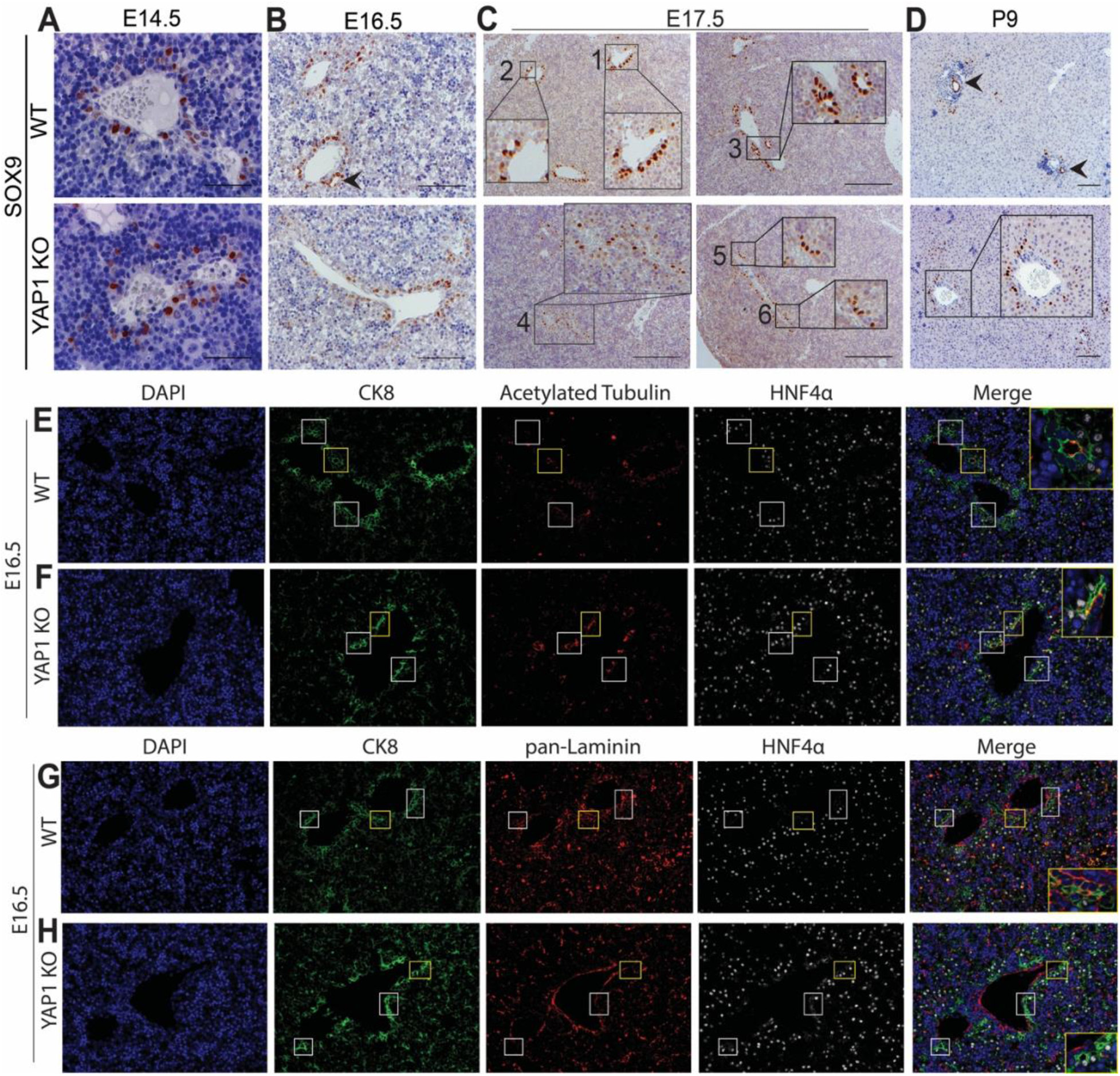
YAP1 KO leads to defective bile duct morphogenesis in late liver development. IHC for SOX9 in WT and YAP1 KO livers at (A) E14.5, (B) E16.5, (C) E17.5, and (D) P9. Arrows and insets point to various stages of bile duct development as described in the text. E-F. IF co-staining for CK8, acetylated tubulin, and HNF4α at E17.5 in WT and YAP1 KO livers showing bile ducts at intermediate stages of maturation (white boxes) with insets highlighted in yellow boxes. G-H. IF co-staining for CK8, pan-laminin, and HNF4α at E17.5 in WT and YAP1 KO livers showing bile ducts at intermediate stages of maturation (white boxes) with insets highlighted in yellow boxes. Scale bars are (A) 50μm and (B-D) 100μm.

Between E16.5 and E17.5, SOX9 staining in WT livers revealed a hierarchical formation of luminal structures, lined first on one side by the above-described pre-existing SOX9-positive ductal plate cells (Fig.4C, inset 2), and on the other side by another layer of SOX9-positive cells thought to be derived from HBs, which lose HNF4α and gain SOX9 to adopt a BEC identity (Fig.4C, inset 3) (Ober and Lemaigre, 2018). At comparable time points, YAP1 KOs showed disparate ductal morphology. At E17.5, we observed formation of only a few luminal structures in KO closest to the largest portal vessels (Fig.4C, insets 5,6), that were lined by SOX9-positive cells *albeit* only on the portal side. Moreover, we failed to observe any real ducts lined completely by the second layer of SOX9-positive cells (Fig.4C). Postnatally, YAP1 KO mice continued to retain a single layer of SOX9+ cells around the portal veins showing no signs of luminal structures (Fig.4D). The parenchymal or the second layer of would-be biliary cells in nascent ducts consistently retained HNF4α in KO, while in WT these cells lose HNF4α and gain SOX9 expression as they mature into BECs (Fig.4E-F). This resulted in an overall increase in the number of HNF4α-positive cells in E17.5 KO livers compared to matched-WT (Fig.S4D-E).

The appearance of luminal biliary structures suggests establishment of polarity which can be detected by IF for acetylated α-tubulin to visualize primary cilia critical for sensing bile flow and regulating numerous growth factor signaling pathways (Mansini et al., 2018). WT cholangiocytes in maturing bile ducts displayed precisely localized acetylated α-tubulin to the apical surface (Fig.4E). In contrast, KO cholangiocytes showed diffuse localization of acetylated α-tubulin (Fig.4F). This suggests an impaired establishment of polarity in maturing portal cholangiocytes in KO mice.

Crosstalk between the ductal plate and the adjacent portal mesenchyme is critical for bile duct morphogenesis (Ober and Lemaigre, 2018). Specifically, portal fibroblasts deposit laminins containing laminin-α1 on the portal side of the ductal plate to provide an initial foundation and later ductal cells deposit laminin-α5 in their surrounding basal lamina (Ober and Lemaigre, 2018; Tanimizu et al., 2012). While WT BECs successfully transitioned to form ducts lined with laminin on both sides (Figure 4G), KO BECs showed only weak to no laminin deposition even on the portal side and failed to robustly deposit laminin around any cholangiocytes, whether HNF4α positive or negative, and the only laminin observed in KO mice was lining the blood vessels (Fig.4H). All together, these data suggest that YAP1 controls the transition and eventually differentiation of the second layer of portal HBs into biliary ductal cells, and absence of YAP1 prevents the second layer of HBs from fully integrating into a maturing ductal structure, leading to absence of the formation of an intrahepatic biliary tree.

## Discussion

By directly knocking out YAP1 from HBs, we addressed the role of YAP1 in the earliest stages of liver development. We show that YAP1 is not necessary for Notch-driven initiation of biliary differentiation in the ductal plate. However, YAP1 loss interferes with establishment of polarity in the ductal plate and also prevents HBs adjacent to this layer from adopting a biliary phenotype and integrating functionally into the nascent ducts to form a patent lumen. Thus, it seems that YAP1 is an essential mediator of BEC maturation and reprogramming of HBs adjacent to the ductal plate (“second layer” HBs), which have adopted a hepatocyte fate but require a switch to adopt a biliary identity for ductal morphogenesis (Ober and Lemaigre, 2018).

There are several broad hypotheses to explain the molecular basis of this defect requiring future studies. First, YAP1 may be involved in cell-cell communication from ductal plate cells that signal to adjacent HBs, which may depend on the proper polarization of the ductal plate cells. This may or may not be occurring downstream of Notch signaling. A previous study suggested that overgrowth of bile ducts due to *Alb*-Cre Nf2-deletion and subsequent YAP1 activation was ablated by *Notch2* deletion, suggesting Notch activity is downstream of YAP1 in bile duct development (Wu et al., 2017). However, *Notch2* deletion in this model did not prevent bile duct formation and these mice exhibited limited injury. Our data, where YAP1 deletion occurs early during biliary differentiation, we suggest Notch activation to be upstream of YAP1 activation in the ductal plate, but downstream of YAP1 signaling in the second layer of HBs, which is a novel finding. In fact, YAP1 has been shown to regulate *Notch2* and *Jagged1* gene expression (Lee et al., 2016; Tschaharganeh et al., 2013). Second, TGFβ signaling originating from the portal mesenchyme is critical for the formation of the second layer during bile duct morphogenesis (Antoniou et al., 2009; Clotman et al., 2005; Lee et al., 2016), and a known driver of hepatocyte-derived biliary regeneration in a model of Alagille syndrome in which Notch signaling was impaired (Schaub et al., 2018). YAP1 may be the downstream effector of Tgfβ signaling in HBs to regulate their fate-switch to second layer of biliary cells transdifferentiation, downregulating HNF4α in second-layer HB (Lee et al., 2016). Third, YAP1 can influence secretion of extracellular matrix components such as laminin α-5 (Gérard et al., 2017; Ober and Lemaigre, 2018), which was deficient around developing ducts in YAP1 KO mice. Interrupting integrin-laminin signaling during bile duct morphogenesis results in defects similar to YAP1 KO (Tanimizu et al., 2012). Some or all of these molecular events may be contributing to the observed phenotypic defect.

We did not detect biliary regeneration through hepatocyte transdifferentiation in our Alagille syndrome-like model, in contrast to other studies (Andersson et al., 2018; Schaub et al., 2018; Walter et al., 2014). Our results thus demonstrate an absolute YAP1 requirement in hepatocytes to undergo transdifferentiation into cholangiocytes. Although other studies have shown YAP1 activation drives expression of biliary markers in hepatocytes and promotes formation of hepatocyte-derived cholangiocarcinoma (Li et al., 2015; Wang et al., 2018; Yimlamai et al., 2014), we provide evidence that without YAP1 the liver cell identity shift cannot occur.

Our model also provides an opportunity to study the relationship between the intrahepatic and extrahepatic bile ducts (EHBDs), which remains poorly understood (Lemaigre, 2020). Foxa3-Cre did not affect *Yap1* expression in EHBDs including gallbladder, all of which formed normally. Using tissue clearing and confocal imaging, we were able to visualize the gallbladder, cystic duct, and perihilar ducts entering the median lobe in both WT and KO mice. Our 2D and 3D imaging showed that the EHBDs extend farther into the median lobe than other liver lobes. We posit that the DR observed in adult KO mice arises from EHBDs responding to the severe cholestatic injury. This response, which is accompanied by fibrosis and inflammation, and shown to be due to signals from cholangiocytes, is primarily associated with large portal vessels in KO mice (Alvaro et al., 2007), and may be an attempt of EHBDs to expand to try and locate and connect with IHBDs, which are lacking in KO. Our model demonstrates that the EHBDs are unable to regenerate IHBDs, similar to previous studies (Schaub et al., 2018), although enabling such process may provide novel therapies in the future.

Despite the severity of liver disease in YAP1 KO mice, they survived long-term by adapting and reprogramming metabolic, synthetic and detoxification functions, while enhancing proliferation and survival signaling. Such adaptations have been reported in other models of liver disease such as Mdr2 KO and combined hepatic Met-EGFR loss (Katzenellenbogen et al., 2007; Paranjpe et al., 2016). YAP1 KO mice completely invert their bile acid transport to overcome the lack of plumbing for bile excretion. While this led to elevated levels of bile acids and bilirubin in the blood, these toxic components were eliminated from hepatic parenchyma thereby reducing hepatocellular injury. Persistently elevated serum total and conjugated bilirubin levels in young children with Alagille syndrome are associated with more severe liver disease and decreased likelihood of spontaneous improvement over time, similar to that seen in our model (Kamath et al., 2010). The same adaptive changes in bilirubin and bile acid transport observed in YAP1 KO mice may be occurring in patients with severe disease and may indicate maximal hepatocyte adaptation in the context of failed biliary regeneration. Thus, YAP1 activation may be an important disease modifier in patients with Alagille syndrome and other biliary disorders requiring further studies. Overall, the surprising capacity of the liver to survive and adapt may be harnessed therapeutically to better understand how to support patients with chronic liver injury.

## Acknowledgments

Funding was provided by 2T32EB001026-16A1 and 1F30DK121393-01A1 to L.M., and 5R01CA204586-05, 1R01DK62277 and Endowed Chair for Experimental Pathology to S.P.M., and by NIH grant 1P30DK120531-01 to Pittsburgh Liver Research Center (for services provided by the Advanced Cell and Tissue Imaging Core). The study was also in part funded by 2P30CA047904-32 to S.W. and the Center for Biological Imaging. We would like to extend our thanks to Dr. Frederic Lemaigre and Dr. Dean Yimlamai for numerous insightful discussions and feedback.

## Author Contributions

Conceptualization: L.M., S.P.M.

Funding Acquisition: L.M., S.P.M., S.W., A.S., A.W.

Investigation: L.M., J.Z., Q.L., T.P-S., K.S., N.J., R.V., S.K., S.H., M.P., S.S., J.T., P.V.B., A.W., A.B.

Methodology: L.M., A.W., S.W., M.P., S.P.M.

Resources: A.S., A.W., S.W.

Supervision: X.M., P.V.B., D.F., T.P-S., P.S., K.N-B., A.B., S.W., S.P.M.

Writing – Original Draft: L.M., S.P.M.

Visualization – L.M., S.P.M.

## STAR Methods

### RESOURCE AVAILABILITY

#### Lead Contact

Further information and requests for resources and reagents should be directed to and will be fulfilled by the Lead Contact, Satdarshan Monga.

#### Materials Availability

Any and all tissue samples and mice described in this paper can be made available under a fully executed Materials Transfer Agreement. Please contact the corresponding author for details.

#### Data and Code Availability

RNA-sequencing data generated in this study are available at Gene Expression Omnibus, Series GSE157777.

### EXPERIMENTAL MODEL AND SUBJECT DETAILS

#### Animal models

C57BL/6 YAP1^fl/fl^ mice (Jackson Labs Stock No. 027929)(Zhang et al., 2010) were bred into C57BL/6 ROSA-stop^fl/fl^-EYFP mice. These mice were then bred into C57BL/6 *Foxa3-Cre* mice described previously(Tan et al., 2008) to create *Foxa3-Cre* YAP1^fl/fl^ ROSA-stop^fl/fl^-EYFP mice (YAP1 KO). Wild type littermate controls were compared to YAP1 KO mice for all subsequent analyses. All animal studies were performed in accordance with the guidelines of the Institutional Animal Use and Care Committee at the University of Pittsburgh School of Medicine and the National Institutes of Health. All animals were group housed in ventilated cages under 12h light/dark cycles with access to enrichment, water and standard chow diet ad libitum unless otherwise specified. Both male and female mice were used throughout the study and littermates were used as WT controls. Mice were analyzed at the following time points: E14.5, E16.5, E17.5, P9, P21, 3-4 months, and 6-8 months of age. Analysis of serum liver function tests was performed by the clinical laboratories at University of Pittsburgh Medical Center (UPMC).

### METHOD DETAILS

#### scRNA-seq Raw Data Retrieval and Processing

The raw sequencing data files were downloaded from public dataset GEO:GSE90047 using the SRA toolkit with the “fastq-dump --split-files” command (http://ncbi.github.io/sra-tools/). The single-end reads were then quantified and aligned using kallisto algorithm with “kallisto quant -i index_file -o output_file -t64 --pseudobam --single -l 51 -s 1 fastq_file”.(Bray et al., 2016) We also used the “samtools sort” command(Li et al., 2009) to sort the aligned bam files before we ran Velocyto(La Manno et al., 2018) to create loom files with the command “velocyto run -e sample_id --onefilepercell --without-umi sorted_bam_file Mus_musculus.GRCm38.100.gtf”. The loom files, which contain the data matrices, were then merged to a single file using the “loompy.combine” function on Python. Finally, the Seurat-Wrappers function “ReadVelocity()” was used to read the combined loom file and convert it to a single-cell gene expression count matrix.(Stuart et al., 2019)

#### scRNA-seq Data Analysis

We used Seurat and Monocle packages to perform the single-cell and the pseudotime analyses.(Stuart et al., 2019; Trapnell et al., 2014) The count matrix was first converted into a Seurat object and then the data was normalized and scaled using Seurat functions NormalizeData() and ScaleData(), respectively. The top 5000 highly variable genes were then selected for the downstream analysis using the Seurat:FindVariableFeatures() function with selection.method = “disp” which selects the genes with the highest dispersion values. The top 5000 highly variable genes were used to perform principal component analysis which was conducted using the Seurat:RunPCA() function. The first 10 principal components were used for Louvain clustering and tSNE visualization (Seurat functions FindClusters() and RunTSNE(), respectively). The expression values of the genes were visualized using Seurat function VlnPlot(). The Pseudotime analysis was performed using the Monocle package following the three standard workflow steps which include choosing genes that define cells’ progress (i.e., feature selection), reducing the data dimensionality using the reverse graph embedding algorithm(Qi et al., 2017), and ordering the cells in pseudotime. The Monocle function plot_genes_in_pseudotime() was used to create Supplemental Figure 1D which shows the expression levels of the genes of interest as a function of the differentiation pseudotime.

#### Immunostaining

Adult livers were harvested and fixed in 10% formalin for 48 hours, then transferred into 70% ethanol followed by paraffin embedding at the UPMC clinical laboratories. Embryonic and fetal livers were harvested and fixed in 4% paraformaldehyde (PFA) for 24 hours at 4C, followed by paraffin embedding. For immunostaining, 4μm paraffin sections were cut, deparaffinized and rehydrated. Sections underwent antigen retrieval by the following methods: 1) pressure cooker, 20 minutes, in sodium citrate buffer pH 6 (YAP1, SOX9, HNF4α, EYFP, CK8, pan-laminin, HES1, CK19 for IF); 2) pressure cooker, 20 minutes, in Agilent DAKO (S1699) target retrieval solution (CK19 for IHC); 3) microwave, 60% power, 12 minutes, in sodium citrate buffer pH 6 (CK8, acetylated tubulin, osteopontin, HNF4α); 4) steamer, 20 minutes, in sodium citrate buffer pH 6 (CD45).

At this point, for immunohistochemistry, slides were treated with 3% H2O2 for 10 minutes to deactivate endogenous peroxidases, washed three times with PBS, then blocked for 10 minutes with SuperBlock reagent (ScyTek Laboratories, AAA500). Slides were incubated in primary antibody diluted in PBS with 0.1% bovine serum albumin (BSA) and 0.01% sodium azide (IHC buffer), for either 1 hour at room temperature (CD45, 1:100) or overnight at 4C (YAP1 1:50, SOX9 1:2000, CK19, 1:50). Slides were then washed three times with PBS and incubated in the appropriate biotinylated secondary antibody at 1:500 dilution for 30 minutes at room temperature. Samples were washed with PBS three times and sensitized with the Vectastain ABC kit (Vector Laboratories, PK-6101) for 30 minutes. Following three washes with PBS color was developed with DAB Peroxidase Substrate Kit (Vector Laboratories, SK-4100), followed by quenching in distilled water for five minutes. Slides were counterstained with hematoxylin (Thermo Scientific, 7211), dehydrated to xylene and coverslips applied with Cytoseal™ XYL (Thermo Scientific, 8312-4). Images were taken on a Zeiss Axioskop 40 inverted brightfield microscope. Whole slides were scanned at 40x magnification using an Aperio AT2 slide scanner (Leica Biosystems).

For immunofluorescence staining, following deparaffinization, rehydration, and antigen retrieval as listed above, sections were permeabilized for 5 minutes with PBS/0.3% TritonX-100 and blocked for 45 minutes in PBS/0.3% TritonX-100/10% BSA. Slides were incubated at 4C overnight in primary antibody cocktails diluted in PBS/0.3% TritonX-100/10% BSA at the following concentrations: CK8, 1:8; all others, 1:100. Slides were washed three times in PBS/0.1% TritonX-100 and incubated at room temperature in secondary antibody cocktails (Invitrogen) also diluted in PBS/0.3% TritonX-100/10% BSA, for 1 hour (dilution 1:500) or 2 hours (dilution 1:800). Slides were again washed three times in PBS/0.1% TritonX-100, then washed three times in PBS, and mounted and coverslipped using VECTASHIELD Antifade Mounting Medium with DAPI (Vector Labs). Slides were imaged on a Nikon Eclipse Ti epifluorescence microscope or LSM 700 Carl Zeiss confocal microscope. Cell and nuclei quantification was performed using Fiji/ImageJ (Schindelin et al., 2012).

For H&E staining, samples were deparaffinized and stained with hematoxylin (Thermo Scientific, 7211) and eosin (Thermo Scientific, 71204), followed by dehydration to xylene and application of a coverslip. For Sirius Red staining, samples were deparaffinized and incubated for one hour in Picro-Sirius Red Stain (American MasterTech, STPSRPT), washed twice in 0.5% acetic acid water, dehydrated to xylene, and coverslipped. Images were taken on a Zeiss Axioskop 40 inverted brightfield microscope.

#### Liver tissue clearing and whole liver immunostaining

Livers were washed in PBS and tissue fixation was achieved by incubating either whole livers or individual liver lobes in 4% paraformaldehyde (PFA) for 24 hours at 4C. Livers were subsequently washed in PBS and stored long-term in PBS/0.1% sodium azide (PBSA) at 4C.

Tissues were incubated in an inactive hydrogel solution overnight at 4C, consisting of 4% acrylamide (Bio-Rad 161-0140), 0.05% bis-acrylamide (Bio-Rad 161-0142), and 0.25% (wt/vol) VA-044 dissolved in PBS. The hydrogel solution was then polymerized by placing the tissues in a water bath at 37C for 3 hours.(Chung et al., 2013; Muntifering et al., 2018) Excess hydrogel was removed, tissues were washed in PBS, and then tissues were placed in the X-CLARITY clearing apparatus (LogosBio C30001). Tissues were cleared using X-CLARITY-ETC Tissue Clearing Solution (LogosBio C13001) supplemented with 10-20mL of N,N,N’,N’-Tetrakis(2-Hydroxypropyl)ethylenediamine (Quadrol) for every 1.5L of clearing solution. The clearing apparatus was run at 37C with constant fluid circulation (100rpm peristaltic pump setting) and a setting of maximum current and voltage set at 1.5 Amps and 70 volts respectively. Timing of tissue clearing varied based on the size of the liver lobes and ranged from 24 hours (smallest lobes, 1000×1000×3mm) to 72 hours (large lobes, 2000×2000×5mm). Next, tissues were washed with PBSA and incubated in 3% H_2_O_2_ for 24 hours. At this stage, after washing with PBSA, tissue could be stored long-term in PBSA at room temperature or 4C, or they could proceed directly to immunostaining.

Tissues were stained as described by Muntifering, *et al* in 2018 with primary and secondary antibodies.(Muntifering et al., 2018) To ensure even staining throughout the sample, antibodies were applied by using the SWITCH protocol.(Murray et al., 2015) All incubations took place at 30C. Briefly, tissues were incubated for 6 days in IHC buffer with 0.5 mM SDS containing CK19 antibody (1:10 dilution). Tissues were then removed from the primary antibody solution and incubated for 1 day in IHC buffer without SDS. Tissues were washed in PBS 3 times for 2 hours each, then incubated in IHC buffer with 0.5mM SDS containing the corresponding conjugated secondary antibody for 6 days. Tissues were then removed from the secondary antibody solution and incubated for 1 day in IHC buffer without SDS. Tissues were washed in PBSA 3 times for 2 hours each, fixed for 2 hours in 4% PFA, washed once more in PBSA, and finally placed in CUBIC R2 solution (50wt% sucrose, 25wt% urea, 10wt% 2,2’,2’’-nitrilotriethanol, and 0.1% (v/v) Triton X-100)(Susaki et al., 2014) for imaging and long-term storage at room temperature.

All tissues were mounted in CUBIC R2 solution and imaged using an RSG4 ribbon scanning confocal microscope (Caliber, Andover, MA) as previously described by Watson *et al*.(Watson et al., 2017) The microscope was fitted with a Nikon CFI90 20x glycerol-immersion objective (Nikon, Melville, NY) with 8.3mm working distance. Volumes were captured with voxel resolution of 0.467 x 0.467 x 12.2 μm (x, y, z). Laser intensity and detector settings were specific to each sample based on the levels of staining. In all cases, the intensity of the laser was increased in a linear manner throughout deeper focal planes to compensate for absorption of excitation and emission light. RAW images acquired in this way were stitched and assembled into composites using a 24 node, 608 core cluster, then converted into the Imaris file format (Bitplane, Zurich, Switzerland). Volumes were rendered using Imaris v9.5.1.

#### RNA extraction and RNA-sequencing analysis

Frozen liver tissue was homogenized in Trizol at 4C and RNA was extracted using QIAGEN RNeasy Mini Kit (Cat. 74104). DNA digestion and removal were performed on the column using the RNase-free DNase Set (Cat. 79254) as per manufacturer instructions. RNA quality and concentration were assessed using a Nanodrop. Purified, high quality RNA from 6 WT livers (3 male, 3 female), and 6 YAP1 KO livers (3 male, 3 female) to Novogene Co. (Sacramento, CA) for cDNA library preparation and RNA-sequencing by Illumina Novaseq 6000 of paired-end 150bp reads, with 20 million reads per end per sample. Raw sequencing data was processed using CLC Genomics Workbench 20.0.3 (QIAGEN Inc., https://digitalinsights.qiagen.com) for quality control and aligned to the *Mus musculus* genome version GRCm38.p6. Reads assigned to each gene underwent TMM normalization and differential expression analysis was performed using *edgeR* within CLC Genomics to compare WT versus YAP1 KO mice. The top differentially expressed genes were filtered by adjusted p-value q < 0.05 and fold-change greater than 2 for subsequent downstream pathway analysis using Ingenuity Pathway Analysis (IPA; QIAGEN Inc., https://www.qiagenbioinformatics.com/products/ingenuity-pathway-analysis), Gene Set Enrichment Analysis (GSEA) and Molecular Signature Database (MSigDB)(Subramanian et al., 2005), and Enrichr(Chen et al., 2013; Kuleshov et al., 2016). CLC Genomics Workbench, and IPA were all used under commercial licenses acquired by the University of Pittsburgh Health Sciences Library System.

#### Measurement of bile flow through cannulation of the common bile duct

The common bile duct was cannulated and the bile flow rate was measured in live three to four month-old, male and female, WT and YAP1 KO mice. The bile duct was cannulated with a microfil tubing according to previously described techniques.(Plaa and Becker, 1965; Tonsberg et al., 2010) In brief, mice were anesthetized with Avertin 0.5 mg/g intraperitoneally (IP). The common bile duct was incised with a pair of fine iridectomy scissors about 6 mm below the hilum of the liver. A microfil tube (WPI, Sarasota, FL, MF28G-5) was passed through the incision and propelled towards the hilum for a distance of about 3 mm. Bile flow rate was recorded (μl/min/100g body weight) and bile was collected in CryoTube vials (Thermo Fisher Scientific) and immediately placed in liquid nitrogen. Animal work described in this manuscript has been approved and conducted under the oversight of the University of Pittsburgh Institutional Animal Care and Use Committee.

#### Bile acid species detection and quantification

Bile acid profiling was performed as described previously.(Zhu et al., 2018) For liver tissue samples, livers were homogenized in water (100 mg tissue in 500 μL water), and then 300μL of methanol: acetonitrile (v/v, 1:1) was added to a 100 μL aliquot of liver homogenate. For serum samples, 25 μL serum was mixed with 100 μL of methanol: acetonitrile (v/v, 1:1). All the mixtures were vortexed for 2 min and centrifuged at 15,000 rpm for 10 minutes. Two microliter of the supernatants from all samples was injected into the ultra-performance liquid chromatography (UPLC) coupled with a SYNAPT G2-S quadrupole time-of-flight mass spectrometry (Waters Corporation, Milford, MA) for analysis. The column type is Acquity UPLC BEH C18 column (2.1 × 100 mm, 1.7 μm). The details of mobile phase gradient were reported previously.(Jiang et al., 2015) The QTOFMS system was operated in a negative high-resolution mode with electrospray ionization as described previously.(Zhu et al., 2018) Bile acid species were quantified by measuring their relative abundance as the area under the curve for each species using standards for comparison. WT and YAP1 KO liver samples or serum samples were compared using a t-test followed by Benjamini-Hochberg correction for multiple hypothesis testing, using FDR < 0.1 as a cutoff for significance.

#### Quantitative liver intravital imaging

Surgical methods used for the intravital imaging were described previously by Pradhan-Sundd, *et al*. (Pradhan-Sundd et al., 2018) Intravascular fluorescent dyes included 100 μg of Carboxyflurescein (CF) and 200 μg of TXR dextran. TXR dextran (MW 70,000) was used to visualize the blood flow through the liver sinusoids whereas CF (MW 377) was used to visualize uptake of the dye from blood to hepatocytes at 1-2m post-injection and then from the hepatocyte to the bile-canaliculi within 5m. Microscopy was performed using a Nikon MPE multi-photon excitation microscope. Movies were processed using Nikon’s NIS Elements (Nikon Elements 3.10). Signal contrast in each channel of a multicolor image was further enhanced by adjusting the maxima and minima of the intensity histogram of that channel. A median filter with a kernel size of 3 was applied over each video frame to improve signal-to-noise ratio.

### QUANTIFICATION AND STATISTICAL ANALYSIS

Statistical details of each experiment can be found in the respective figure legends. Data are presented as mean ± standard deviation (sd). n refers to biological replicates. p < 0.05 was considered statistically significant, except for individual bile acid species comparisons for which an FDR of 0.1 was used. GraphPad PRISM 7.0c software was used for statistical analyses. No samples or animals were excluded from the analysis. Littermate controls of both genders were used throughout the study. Data significance was analyzed using a two-tailed unpaired Student’s t test or Mann-Whitney test in cases where two groups were being compared. In cases where more than two groups were being compared, one-way or two-way ANOVA were used with Sidak’s test to correct for multiple comparisons.

## KEY RESOURCES TABLE

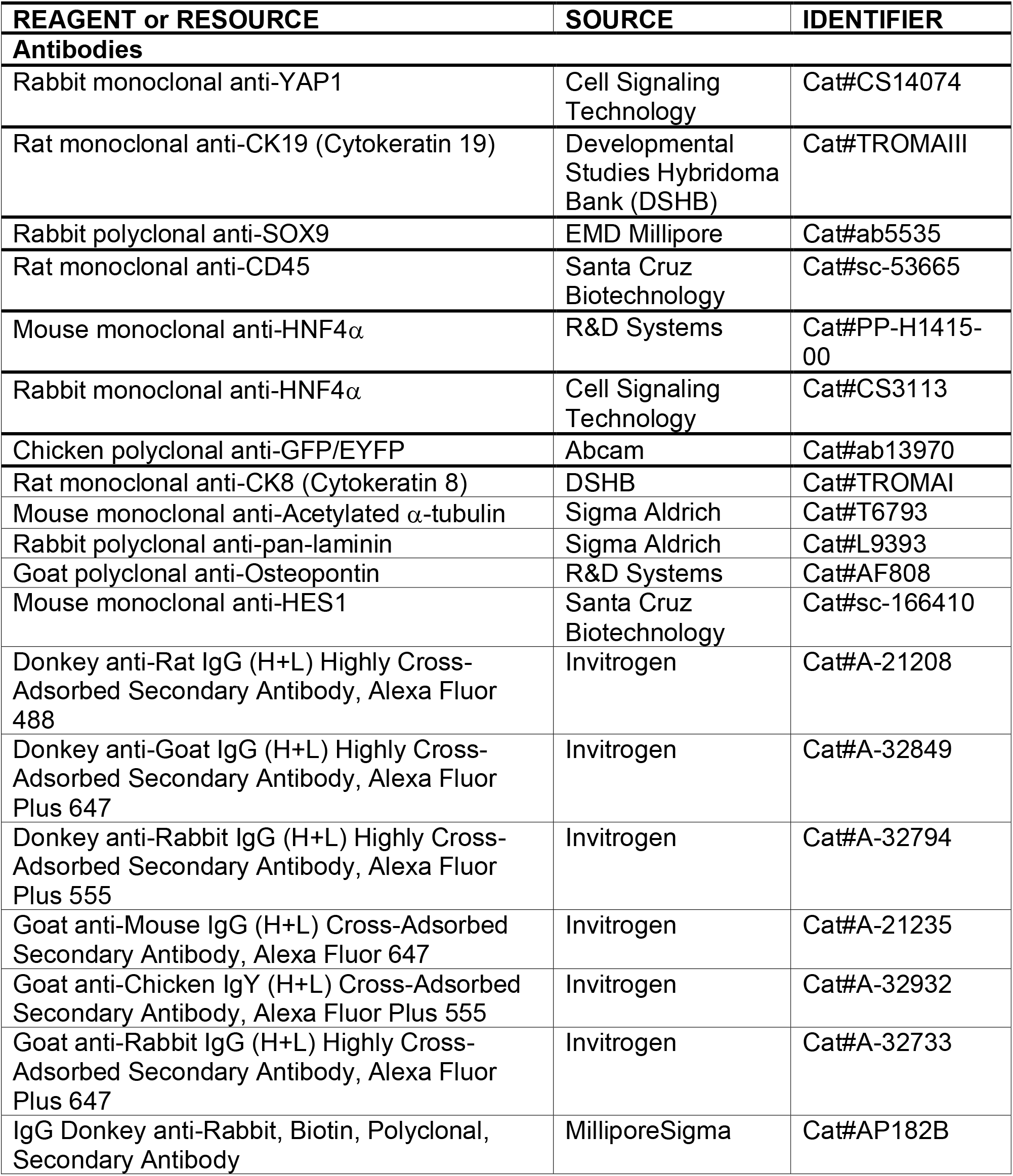
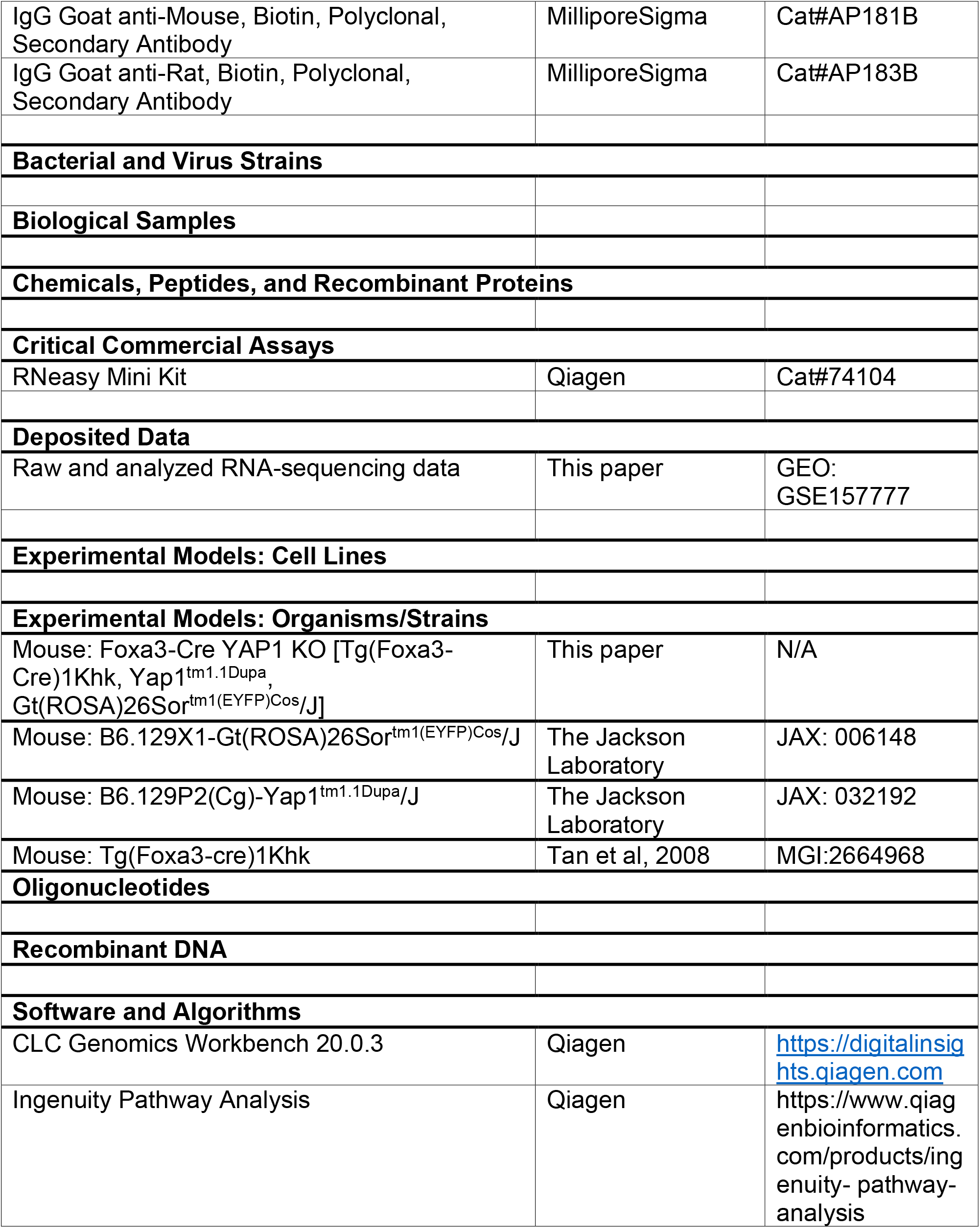
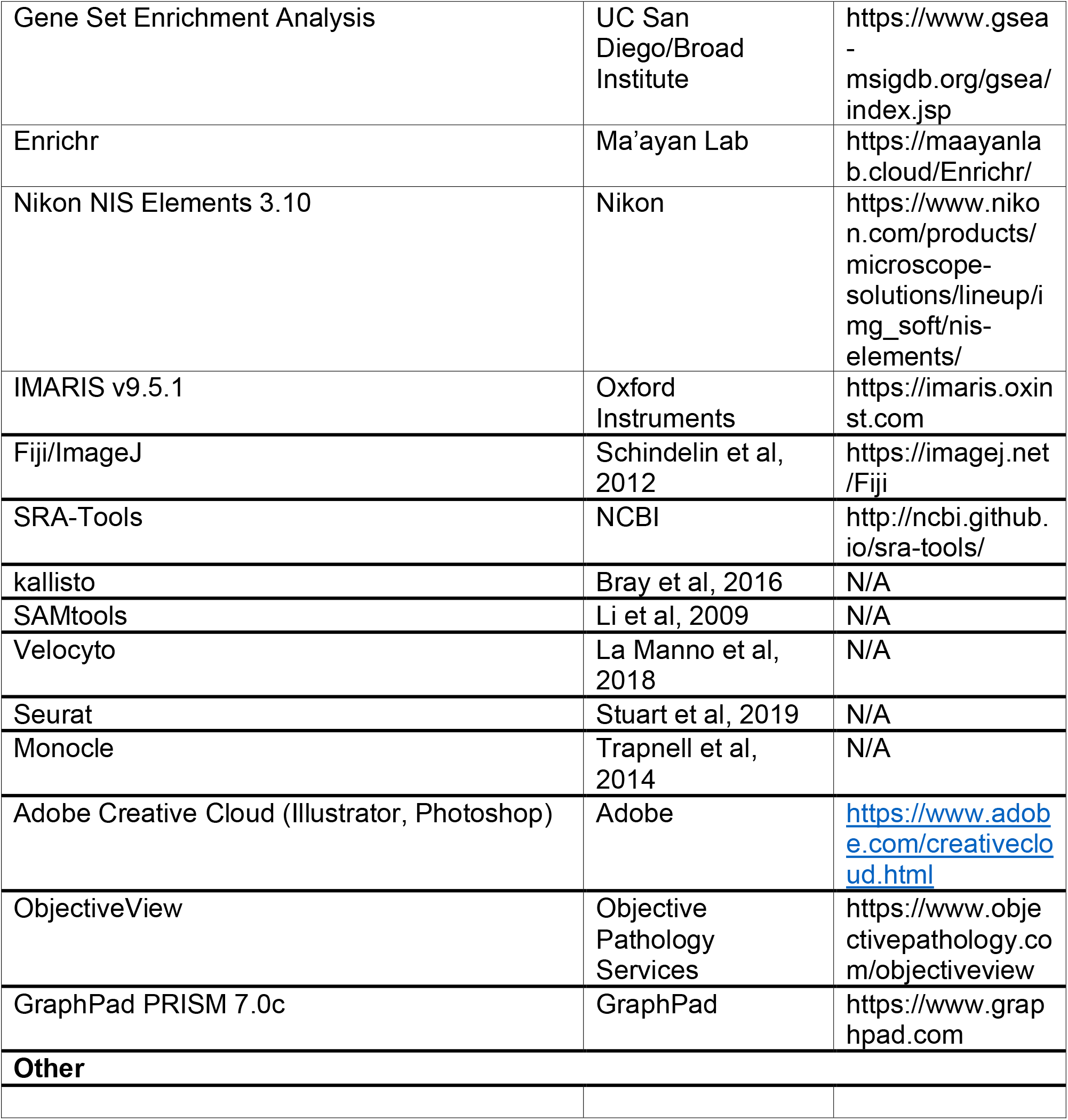

## Supplemental Information Titles and Legends

**Supplemental Figure 1. Characterization of YAP1 activity in early liver development and phenotype of YAP1 KO mice.**

A-B. tSNE plots of single cell sequencing data from a previously published study (GSE90047) on embryonic liver epithelial cells reveals three clusters (a, b, and c) of cells roughly compatible with the original publication’s classification of hepatoblasts, hepatocytes, and cholangiocytes (Cell Type) and traces the divergence of both lineages over pseudotime and developmental time from E10.5 to E17.5. Boxes highlight cells which cluster with hepatoblasts but were identified as cholangiocytes and are labeled as cluster d in panels C and D. C. Gene expression of key cell type markers and YAP1 targets by cluster. D. Gene expression of key cell type markers and YAP1 targets by cell type and pseudotime. E. IF co-staining for HNF4α and YAP1 at E14.5 in WT and YAP1 KO liver. F. H&E shows patches of necrosis in KO mice compared to healthy WT. G. SOX9 IHC shows clusters of SOX9+ hepatocytes in contrast to mature bile ducts in WT. H-I. KO mice show increased fibrosis (Sirius Red) and inflammation (CD45) vs WT, particularly around areas of ductular reaction. Scale bars (F-I) are 100μm.

**Supplemental Figure 2. Pathway analysis of RNA-sequencing data comparing adult YAP1 KO mice to WT.**

A. Principal component analysis of RNA-sequencing data clearly distinguishes between WT and KO mice. B. IPA analysis highlighted several altered pathways related to liver fibrosis in YAP1 KO vs WT. C-D. GSEA showed increased enrichment of numerous inflammatory and cancer-related pathways in KO vs WT mice. Enrichr analysis of regulatory regions common to genes upregulated (E) or downregulated (F) in KO mice vs WT identified key transcription factors whose activity is altered in KO mice.

**Supplemental Figure 3. Quantification of bile acid species by mass spectrometry in adult liver tissue and serum of both male and female WT and YAP1 KO mice.**

A-C. Primary unconjugated (A), primary conjugated (B), and secondary (C) bile acids in liver tissue of male mice. D-F. Primary unconjugated (D), primary conjugated (E), and secondary (F) bile acids in serum of male mice. G-I. Primary unconjugated (G), primary conjugated (H), and secondary (I) bile acids in liver tissue of female mice. J-L. Primary unconjugated (J), primary conjugated (K), and secondary (L) bile acids in serum of female mice. WT vs KO values for each species by gender and sample type were analyzed by t-test, and all p-values were adjusted by Benjamini-Hochberg correction for multiple hypothesis testing, with an FDR of 0.1 (*p< 0.05, **p < 0.001). Data show mean ± sd.

**Supplemental Figure 4. Characterization of bile duct maturation in embryonic development of YAP1 KO mice.**

A. IF co-staining for EYFP and HNF4α in YAP1 KO mice. B. Quantification of EYFP labeling of HNF4α+ hepatoblasts at E17.5 in KO mice. C. IHC for HES1 showing expression in ductal plate of WT and KO mice at E16.5. D. IF staining for HNF4α+ in WT and KO mice at E17.5. E. Quantification of HNF4α+ nuclei in KO vs WT mice (t-test, * p< 0.05).

**Supplemental Movie 1. 3D imaging of WT and YAP1 KO biliary tree**

Whole tissue clearing, IF for CK19, and ribbon scanning confocal microscopy were used to visualize the biliary tree in three-dimensions. Here we show the 3D representations of the biliary tree in A) WT mouse at P21, median lobe and gallbladder; B) YAP1 KO mouse at P21, median lobe and gallbladder; C) WT mouse at 8 months, left liver lobe; D) YAP1 KO mouse at 8 months, left liver lobe.

**Supplemental Movie 2. Intravital imaging of adult WT and YAP1 KO mice**

Intravital imaging was used to trace the flow of blood (red) and bile (green) in A) WT and B) YAP1 KO mice at 3-4 months of age.

