## Supplemental Figures for "Compensatory hepatic adaptation accompanies permanent absence of intrahepatic biliary network due to YAP1 loss in liver progenitors"

**Supplemental Figures and Legends**

1. **4 figures (S1-4)**
2. **2 movies (Movie 1-2)**

**Figure S1**

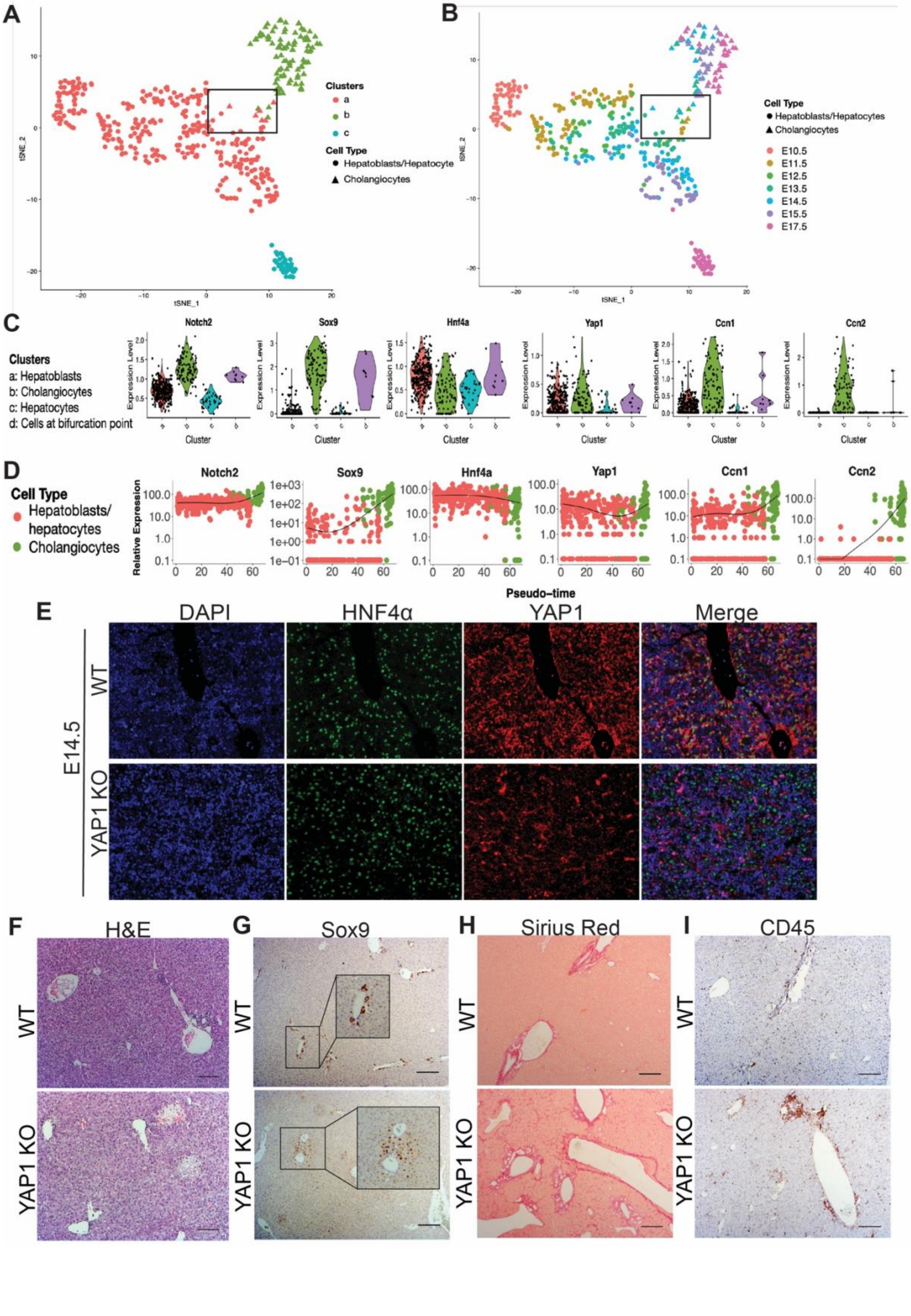
A
B
Clusters
a
b
c
Cell Type
Hepatoblasts/Hepatocyte
Cholangiocytes
tSNE_1
tSNE_2
Cell Type
Hepatoblasts/Hepatocytes
Cholangiocytes
E10.5
E11.5
E12.5
E13.5
E14.5
E15.5
E17.5
C
Clusters
a: Hepatoblasts
b: Cholangiocytes
c: Hepatocytes
d: Cells at bifurcation point
Notch2
Sox9
Hnf4a
Yap1
Ccn1
Ccn2
Expression Level
Cluster
D
Cell Type
Hepatoblasts/hepatocytes
Cholangiocytes
Relative Expression
Pseudo-time
E
DAPI
HNF4α
YAP1
Merge
E14.5
WT
YAP1 KO
F
H&E
G
Sox9
H
Sirius Red
I
CD45
WT
YAP1 KO

**Figure S1. Characterization of YAP1 activity in early liver development and phenotype of YAP1 KO mice.**

A-B. tSNE plots of single cell sequencing data from a previously published study (GSE90047) on embryonic liver epithelial cells reveals three clusters (a, b, and c) of cells roughly compatible with the original publication's classification of hepatoblasts, hepatocytes, and cholangiocytes (Cell Type) and traces the divergence of both lineages over pseudotime and developmental time from E10.5 to E17.5. Boxes highlight cells which cluster with hepatoblasts but were identified as cholangiocytes and are labeled as cluster d in panels C and D. C. Gene expression of key cell type markers and YAP1 targets by cluster. D. Gene expression of key cell type markers and YAP1 targets by cell type and pseudotime. E. IF co-staining for HNF4$\alpha$ and YAP1 at E14.5 in WT and YAP1 KO liver. F. H&E shows patches of necrosis in KO mice compared to healthy WT. G. SOX9 IHC shows clusters of SOX9+ hepatocytes in contrast to mature bile ducts in WT. H-I. KO mice show increased fibrosis (Sirius Red) and inflammation (CD45) vs WT, particularly around areas of ductular reaction. Scale bars (F-I) are 100$\mu$m.

**Figure S2**

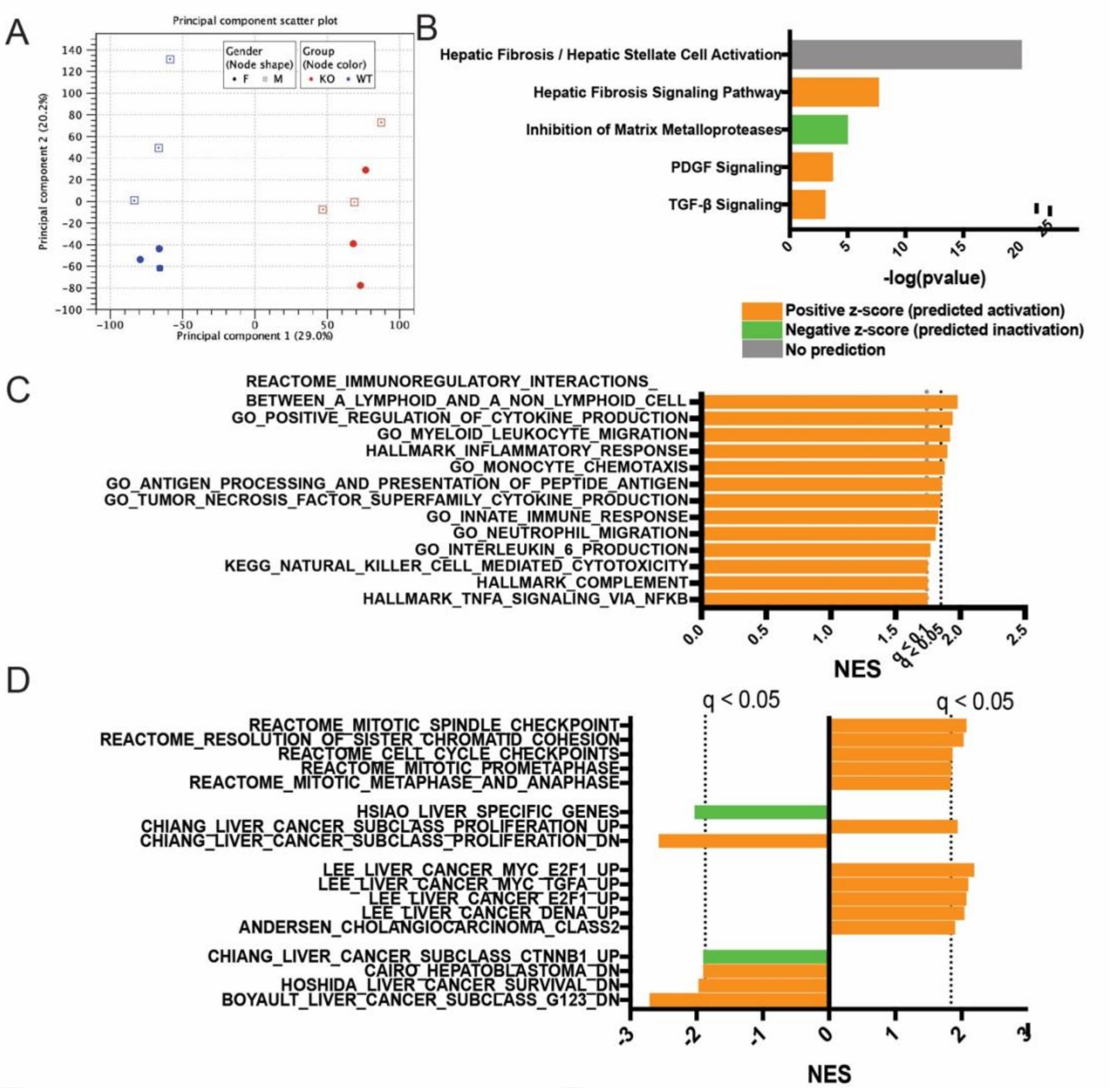


E

| ENCODE and ChEA TF Consensus Regions - Enriched among regulatory regions of genes upregulated in YAP1 KO mice | | | | | |
|---|---|---|---|---|---|
| Index | Name | P-value | Adjusted p-value | Odds Ratio | Combined score |
| 1 | FOXM1 ENCODE | 3.45E-13 | 1.79E-11 | 4.06 | 116.37 |
| 2 | SUZ12 CHEA | 2.43E-13 | 2.53E-11 | 1.58 | 45.93 |
| 3 | SPI1 CHEA | 1.13E-10 | 3.91E-09 | 1.66 | 38.1 |
| 4 | IRF8 CHEA | 6.66E-10 | 1.73E-08 | 3.18 | 67.28 |
| 5 | E2F4 ENCODE | 3.31E-09 | 6.88E-08 | 1.76 | 34.28 |
| 6 | RUNX1 CHEA | 4.43E-09 | 7.68E-08 | 1.53 | 29.48 |
| 7 | TP63 CHEA | 4.08E-07 | 6.1E-06 | 1.43 | 21.09 |
| 8 | TP53 CHEA | 1.1E-05 | 0.00015 | 1.85 | 21.06 |
| 9 | NFE2L2 CHEA | 4.4E-05 | 0.00051 | 1.41 | 14.13 |
| 10 | STAT3 CHEA | 7.3E-05 | 0.00075 | 2.05 | 19.53 |
| 11 | SPI1 ENCODE | 0.00045 | 0.00428 | 1.27 | 9.8 |
| 12 | SMAD4 CHEA | 0.00062 | 0.00535 | 1.46 | 10.75 |
| 13 | PPARG CHEA | 0.00142 | 0.01133 | 1.44 | 9.45 |
| 14 | SALL4 CHEA | 0.00197 | 0.01461 | 1.53 | 9.55 |
| 15 | VDR CHEA | 0.00236 | 0.01638 | 2.24 | 13.53 |
| 16 | GATA2 CHEA | 0.00379 | 0.02464 | 1.32 | 7.36 |
| 17 | FOSL2 ENCODE | 0.00404 | 0.02472 | 1.68 | 9.24 |
| 18 | GATA1 CHEA | 0.00456 | 0.02632 | 1.31 | 7.04 |

F

| ChEA 2016 TF Regulators - Enriched among regulatory regions of genes downregulated in YAP1 KO mice | | | | | |
|---|---|---|---|---|---|
| Index | Name | P-value | Adjusted p-value | Odds Ratio | Combined score |
| 1 | LXR 22158963 ChIP-Seq LIVER Mouse | 5.05E-23 | 3.26E-20 | 2.41 | 123.8 |
| 2 | RXR 22158963 ChIP-Seq LIVER Mouse | 4.50E-22 | 1.45E-19 | 2.38 | 116.78 |
| 3 | PPARA 22158963 ChIP-Seq LIVER Mouse | 5.89E-14 | 1.27E-11 | 2.04 | 62.12 |
| 4 | ESR1 17901129 ChIP-ChIP LIVER Mouse | 1.53E-08 | 2.5E-06 | 2.88 | 51.73 |
| 5 | FOXO1 23066095 ChIP-Seq LIVER Mouse | 6.85E-07 | 8.8E-05 | 2.86 | 40.61 |
| 6 | SUZ12 20075857 ChIP-Seq MESCs Mouse | 1.5E-05 | 0.00138 | 1.34 | 14.92 |
| 7 | EGR1 23403033 ChIP-Seq LIVER Mouse | 1.5E-05 | 0.00159 | 1.88 | 20.9 |
| 8 | CLOCK 20551151 ChIP-Seq 293T Human | 0.00025 | 0.01995 | 2.18 | 18.09 |
| 9 | EED 16625203 ChIP-ChIP MESCs Mouse | 0.00038 | 0.0245 | 1.75 | 13.8 |
| 10 | PIAS1 25552417 ChIP-Seq VCAP Human | 0.00037 | 0.02665 | 1.8 | 14.21 |
| 11 | JARID2 20075857 ChIP-Seq MESCs Mouse | 0.00083 | 0.04845 | 1.55 | 11.01 |
| 12 | FOXA2 19822575 ChIP-Seq HepG2 Human | 0.00093 | 0.04994 | 1.33 | 9.26 |

**Figure S2. Pathway analysis of RNA-sequencing data comparing adult YAP1 KO mice to WT.**

A. Principal component analysis of RNA-sequencing data clearly distinguishes between WT and KO mice. B. IPA analysis highlighted several altered pathways related to liver fibrosis in YAP1 KO vs WT. C-D. GSEA showed increased enrichment of numerous inflammatory and cancer-related pathways in KO vs WT mice. Enrichr analysis of regulatory regions common to genes upregulated (E) or downregulated (F) in KO mice vs WT identified key transcription factors whose activity is altered in KO mice.

**Figure S3**

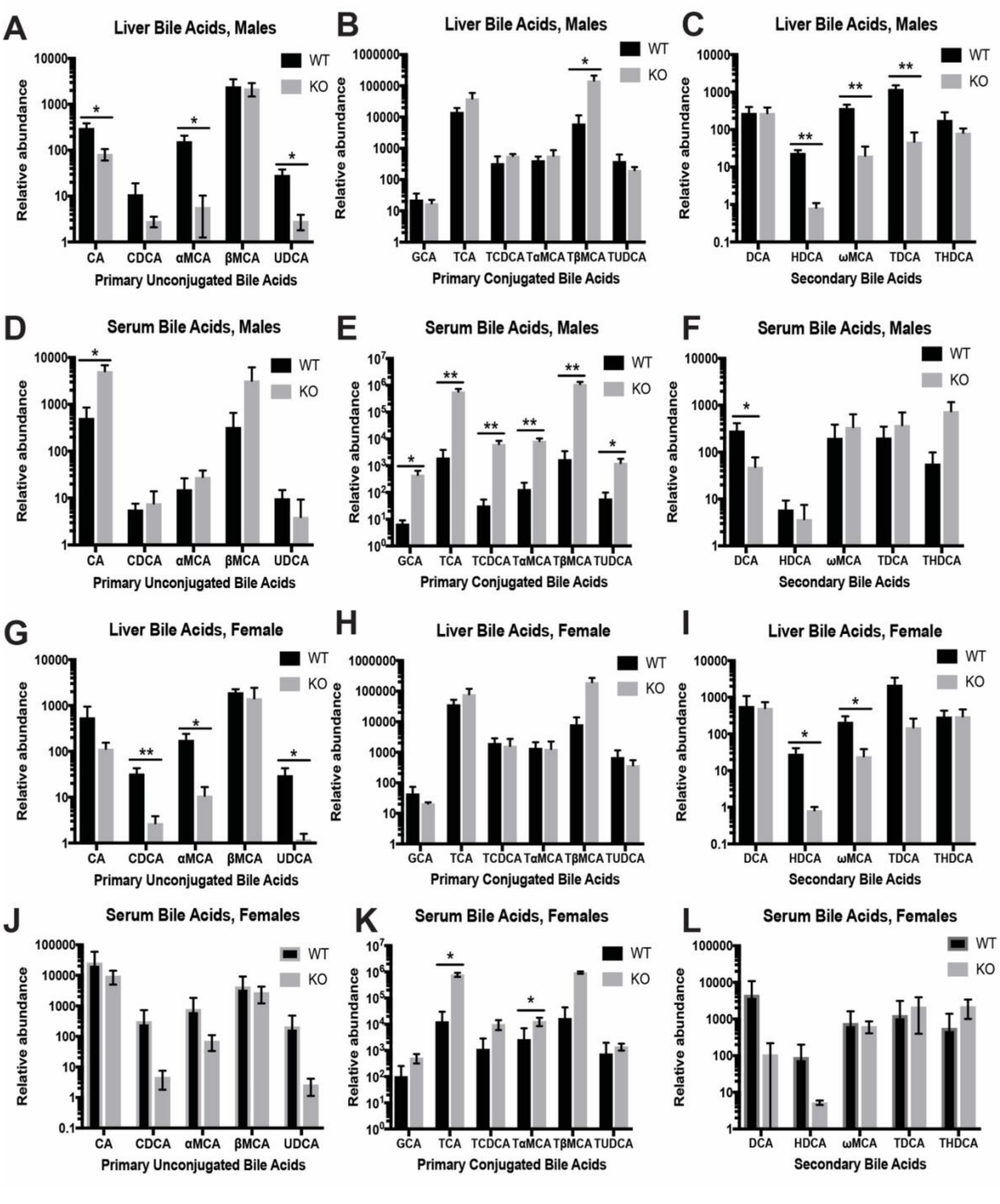
A
Liver Bile Acids, Males
Relative abundance
CA CDCA αMCA βMCA UDCA
Primary Unconjugated Bile Acids
B
Liver Bile Acids, Males
GCA TCA TCDCA TαMCA TβMCA TUDCA
Primary Conjugated Bile Acids
C
Liver Bile Acids, Males
DCA HDCA ωMCA TDCA THDCA
Secondary Bile Acids
D
Serum Bile Acids, Males
E
Serum Bile Acids, Males
F
Serum Bile Acids, Males
G
Liver Bile Acids, Female
H
Liver Bile Acids, Female
I
Liver Bile Acids, Female
J
Serum Bile Acids, Females
K
Serum Bile Acids, Females
L
Serum Bile Acids, Females
WT
KO

**Figure S3. Quantification of bile acid species by mass spectrometry in adult liver tissue and serum of both male and female WT and YAP1 KO mice.**
A-C. Primary unconjugated (A), primary conjugated (B), and secondary (C) bile acids in liver tissue of male mice. D-F. Primary unconjugated (D), primary conjugated (E), and secondary (F) bile acids in serum of male mice. G-I. Primary unconjugated (G), primary conjugated (H), and secondary (I) bile acids in liver tissue of female mice. J-L. Primary unconjugated (J), primary conjugated (K), and secondary (L) bile acids in serum of female mice. WT vs KO values for each species by gender and sample type were analyzed by t-test, and all p-values were adjusted by Benjamini-Hochberg correction for multiple hypothesis testing, with an FDR of 0.1 (*$p < 0.05$, **$p < 0.001$). Data show mean $\pm$ sd.

**Figure S4**

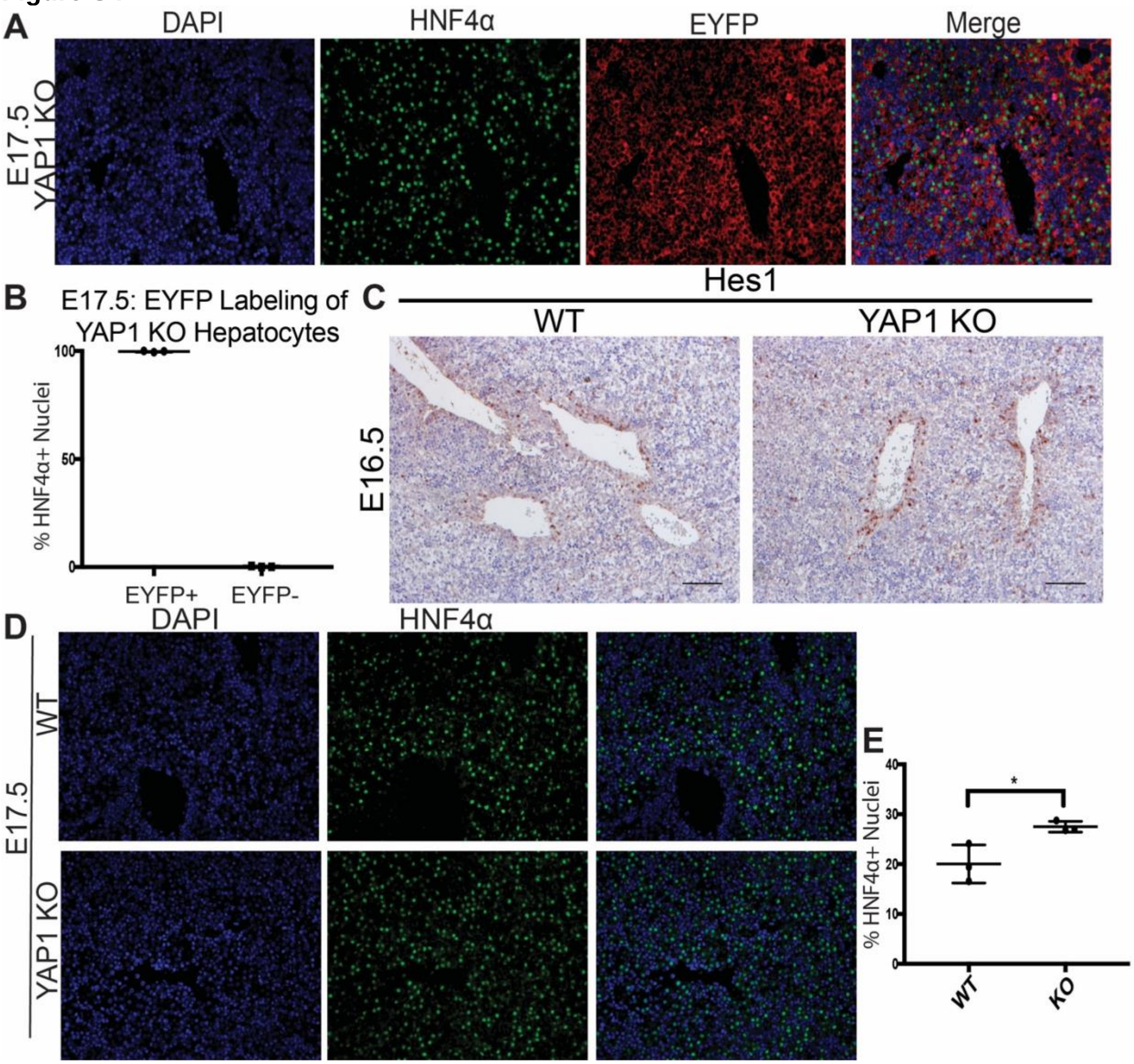


**Figure S4. Characterization of bile duct maturation in embryonic development of YAP1 KO mice.**

A. IF co-staining for EYFP and HNF4α in YAP1 KO mice. B. Quantification of EYFP labeling of HNF4α+ hepatoblasts at E17.5 in KO mice. C. IHC for HES1 showing expression in ductal plate of WT and KO mice at E16.5. D. IF staining for HNF4α+ in WT and KO mice at E17.5. E. Quantification of HNF4α+ nuclei in KO vs WT mice (t-test, * $p < 0.05$).
